# Xylazine Exacerbates Fentanyl-Induced Respiratory Depression and Bradycardia

**DOI:** 10.1101/2024.08.16.608310

**Authors:** Catherine Demery-Poulos, Sierra C. Moore, Erica S. Levitt, Jessica P. Anand, John R. Traynor

## Abstract

Fatal opioid overdoses in the United States have nearly tripled during the past decade, with greater than 92% involving a synthetic opioid like fentanyl. Fentanyl potently activates the μ-opioid receptor to induce both analgesia and respiratory depression. The danger of illicit fentanyl has recently been exacerbated by adulteration with xylazine, an α2-adrenergic receptor agonist typically used as a veterinary anesthetic. In 2023, over a 1,000% increase in xylazine-positive overdoses was reported in some regions of the U.S. Xylazine has been shown to potentiate the lethality of fentanyl in mice, yet a mechanistic underpinning for this effect has not been defined. Herein, we evaluate fentanyl, xylazine, and their combination in whole-body plethysmography (to measure respiration) and pulse oximetry (to measure blood oxygen saturation and heart rate) in male and female CD-1 mice. We show that xylazine decreases breathing rate more than fentanyl by increasing the expiration time. In contrast, fentanyl primarily reduces breathing by inhibiting inspiration, and xylazine exacerbates these effects. Fentanyl but not xylazine decreased blood oxygen saturation, and when combined, xylazine did not change the maximum level of fentanyl-induced hypoxia. Xylazine also reduced heart rate more than fentanyl. Finally, loss in blood oxygen saturation correlated with the frequency of fentanyl-induced apneas, but not breathing rate. Together, these findings provide insight into how the addition of xylazine to illicit fentanyl may increase the risk of overdose.

**Graphical Abstract:** 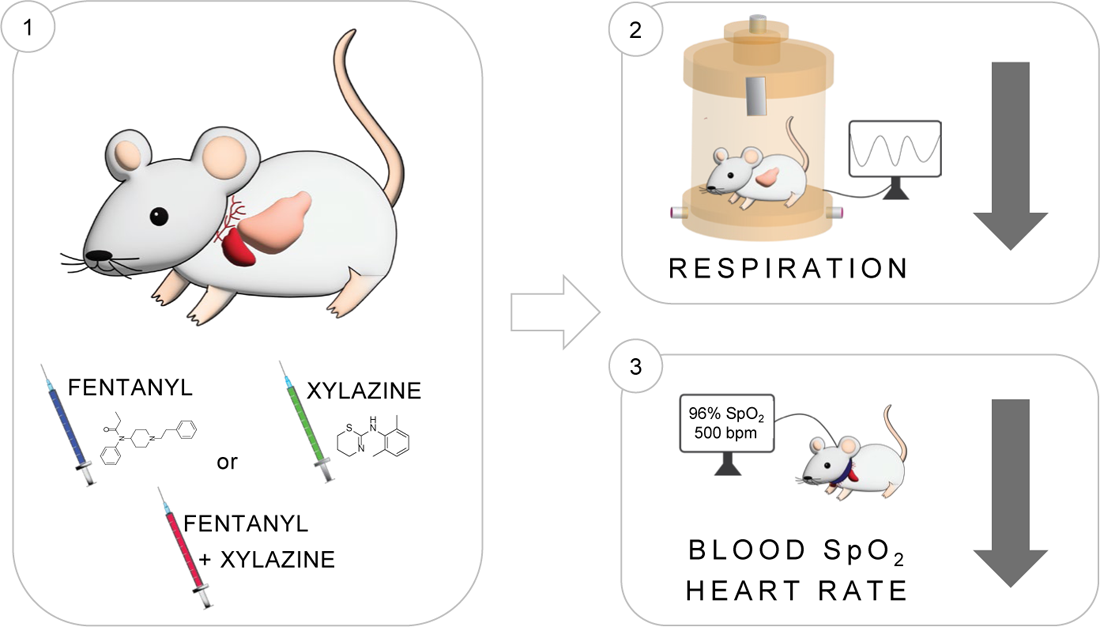

## INTRODUCTION

Fatal opioid overdoses in the United States have drastically increased over the past decade, exceeding 82,000 in 2023. Over 92% of these overdoses involve a synthetic opioid such as fentanyl [1]. Fentanyl, along with other illicit and clinically important opioids, binds to the μ-opioid receptor (MOR) to induce analgesia as well as the life-threatening effect of respiratory depression. Fentanyl is more dangerous than classical opioids like morphine for reasons including greater potency at MOR, increased lipophilicity that facilitates rapid blood-brain barrier penetration, and induction of respiratory muscle rigidity [2–5]. The danger of illicit fentanyl has recently been exacerbated by the replacement of less harmful adulterants (i.e., mannitol) with xylazine, an α2-adrenergic receptor (α2AR) agonist used as a veterinary anesthetic [6–8]. Activation of α2AR autoreceptors leads to central nervous system depression that can increase the likelihood of overdose when combined with opioids. Furthermore, xylazine is not used in humans due to its induction of severe bradycardia [7,9]. The co-detection of fentanyl with xylazine has risen starkly over the past few years, both in DEA seizures [10] and postmortem toxicology reports [11], leading the White House to declare fentanyl-xylazine combinations an emerging threat to the United States [12].

Xylazine is rarely abused independently of opioids, with over 99% of xylazine detection occurring in the presence of fentanyl [8]. Yet, 89% of illicit opioid users who are aware of potential xylazine adulteration do not want xylazine in their fentanyl samples [13], likely due to the risk of xylazine-induced skin ulcers, which are hypothesized to result from peripheral α2AR activation [7]. Information regarding the ratio of fentanyl-to-xylazine in illicit samples is lacking, but postmortem toxicology studies have reported an approximate 1:1 median ratio [14] and a 1:4 mean ratio [15] of fentanyl to xylazine. In mice, xylazine has previously been shown to potentiate the lethality of fentanyl at ratios of 1:5 and 1:10 (fentanyl to xylazine) [16]. Furthermore, isobolographic analysis of fentanyl- and xylazine-induced lethality in mice showed that the two drugs synergize when combined, and that naloxone (an opioid receptor antagonist) but not yohimbine (an α2AR antagonist) was able to reverse the lethal effect of a fentanyl-xylazine combination [17]. Xylazine was also shown to prolong fentanyl-induced brain hypoxia without changing the maximum level of hypoxia [18], an effect that was fully rescued by co-administration of naloxone together with the α2AR antagonist, atipamezole.[19]. Finally, other preclinical studies have found that xylazine does not increase the rewarding effects of fentanyl but can alter symptoms of withdrawal [16,20,21].

Since the lethal effects of fentanyl are due to respiratory depression, in this study we investigated whether xylazine alters fentanyl-induced respiratory depression. We considered which components of the respiratory process were affected using whole-body plethysmography and vital signs using pulse oximetry in male and female CD-1 mice. Fentanyl was held constant at a dose (3.2 mg/kg) that produces significant but not maximal respiratory depression and combined with xylazine at ratios of 1:1 up to 1:10, to best reflect the relative quantities of both drugs identified in human overdose postmortem samples. Changes in breathing rate, inspiration, expiration, apneas, blood oxygen saturation, and heart rate were evaluated. We found that xylazine reduced breathing rate and heart rate more than fentanyl and drove the effect when the two drugs were combined. As expected, fentanyl decreased blood oxygen levels, whereas xylazine had no effect despite reducing breathing rate to a greater extent than fentanyl. Reductions in blood oxygen levels correlated with the frequency of drug-induced apneas. These findings shed light on the mechanisms by which the addition of xylazine to illicit fentanyl may increase the risk of overdose in humans.

## METHODS

### Animal Care and Use

Male and female CD-1 mice aged 6-12 weeks were used for all experiments (n=84 total). Mice were either bred in-house at the University of Michigan or purchased from Envigo (Indianapolis, IN). All experiments were performed during the light phase of the 12-hour light/dark cycle on which mice were maintained. Mice were housed by sex in groups of 2-5 in clear polypropylene home cages with corncob bedding and free access to food, water, and enrichment. All studies followed the Guide for the Care and Use of Laboratory Animals [22] and were approved by the University of Michigan Institutional Animal Care and Use Committee.

### Drug Preparation

Fentanyl powder (Cayman Chemical Company, Ann Arbor, MI) was dissolved in saline. Xylazine (Cayman Chemical Company) powder was dissolved in DMSO, which was then diluted in castor oil followed by saline to achieve a 1:1:8 ratio of DMSO, castor oil, and saline, respectively. For the combined doses of fentanyl + xylazine, fentanyl was prepared in the saline component of the diluent and added last to the preparation of xylazine in DMSO and castor oil to achieve the desired drug concentrations. Vehicle contained 1:1:8 proportions of DMSO, castor oil, and saline, respectively. Each animal received only one injection at 10 mL/kg of bodyweight.

### Whole-Body Plethysmography

Respiratory measures were recorded using vivoFlow^®^ whole-body plethysmography (WBP) chambers by SCIREQ (Montreal, QC Canada). Chambers were supplied with 0.5 L/min of medical grade air USP (Cryogenic Gases). All mice were given 30 min to acclimate to WBP chambers the day prior to test day. On test day, weights and temperatures (Braun digital infrared thermometer) for all mice were recorded. Body temperature, chamber temperature and chamber humidity were used in volume calculations. Awake, free-moving mice were then placed in chambers for 10 min of habituation (data not saved) and 20 min of baseline recording prior to intraperitoneal injection of xylazine, fentanyl + xylazine, or vehicle. Mice were returned to WBP chambers for 60 min of post-treatment recording. Emka Technologies iox 2.10.8.6 software (Sterling, VA) was used to collect data in 30-sec intervals, which were then collapsed into 5-min bins for analysis. Apneas and EF_50_ values were calculated after exporting data as individual breaths. The same vehicle group is shown in xylazine and fentanyl plus xylazine time course graphs.

### Pulse Oximetry

Blood oxygen saturation and heart rate were measured using the MouseOx^®^ Plus Small Animal Pulse Oximeter from Starr Life Sciences (Oakmont, PA). Awake, free-moving animals were habituated to enclosures and practice collars for 60 min prior to experiment start. After, practice collars were exchanged for the MouseOx^®^ collar and baseline recordings were taken for 60 min before mice received an intraperitoneal injection of xylazine, fentanyl plus xylazine, or vehicle. Data was collected for 60 min post-injection and binned into 5-min bins for analysis. The same vehicle group is shown in xylazine and fentanyl plus xylazine time course graphs.

### Data Analysis

For breathing rate, inspiration (Ti) and expiration time (Te), peak inspiratory (PIF) and expiratory flow (PEF), blood oxygen saturation, and heart rate, data were averaged into 5-min bins for analysis. Data are presented as % baseline of each mouse during the 20-min period prior to injection (t=-20-0 min). Raw baseline data by treatment group are shown in **Supp. Table 1**; group averages were compared via one-way ANOVA. All 8 groups were compared from t=0-60 min post-injection via 2-way ANOVA. If the main effect of treatment was p<0.05, average group values from t=10-15 min were then compared via 1-way ANOVA with Tukey’s post-hoc test (p-values displayed in figures) if the main effect was significant with p<0.05. GraphPad Prism 10 was used for all statistical analyses.

For apneas and EF_50_, data were exported as individual breaths for groups that had both plethysmography and pulse oximetry data. Frequency of apneas were calculated in Excel using the following criteria: *Apnea == Te > 2lAvg. Te post-injectionj; Severe Apnea == Te > 2lAvg. Ti + Te post-injectionj*. The average lengths of Ti and Te used to define an apnea were calculated for each mouse individually. Outliers were identified via robust nonlinear regression and outlier removal (ROUT) method (Q=1%) and removed [23]. The average number of apneas per treatment group from t=0-15 min post-injection were compared via 1-way ANOVA with Tukey’s post-hoc test if the main effect was significant with p<0.05. Pearson correlation between blood oxygen saturation (t=10-15 min) and total or severe apneas (t=0-15 min) was computed. EF_50_ values were calculated as % baseline of each mouse during the baseline period (t=-20-0 min). Average EF_50_ values among groups were compared from t=0-15 min or t=0-60 min post-injection using 1-way ANOVA with Tukey’s post-hoc test if the main effect was significant with p<0.05. GraphPad Prism 10 was used for all statistical analyses except linear trendlines and S.E.M. for correlation analyses, which were calculated in Excel.

## RESULTS

### Xylazine reduces breathing rate to a greater degree than fentanyl and drives the effect when the two drugs are combined

We first considered how fentanyl, xylazine, and their combination affected breathing rate using whole-body plethysmography (**Fig. 1A**). The dose of 3.2 mg/kg fentanyl was chosen because it produces significant but non-maximal effects on respiration. Xylazine was combined at a 1:1 up to 10:1 ratio with fentanyl. Given the rapid onset of fentanyl-induced respiratory depression, we compared the average respiration rate of the different treatment groups from 10-15 min post injection (**Fig. 1B** bar graph). All doses of xylazine (3.2, 10, and 32 mg/kg) drastically reduced breathing rate relative to vehicle (p<0.0001). Fentanyl (3.2 mg/kg) also reduced breathing rate relative to vehicle (p<0.0001) but to a lesser extent than the 32 mg/kg dose of xylazine (p<0.05). The combination of fentanyl and 3.2, 10, or 32 mg/kg xylazine further reduced breathing rate relative to fentanyl alone (p<0.01, p<0.0001, and p<0.0001, respectively) but did not significantly differ from the corresponding dose of xylazine alone. In other words, when fentanyl and xylazine are combined, the extent of breathing rate depression is due to xylazine (**Fig 1B**). However, as shown in **Fig. 1C**, fentanyl, xylazine, and their combination induce different breathing patterns that are not captured by the breathing rate metric.

**Figure 1.**
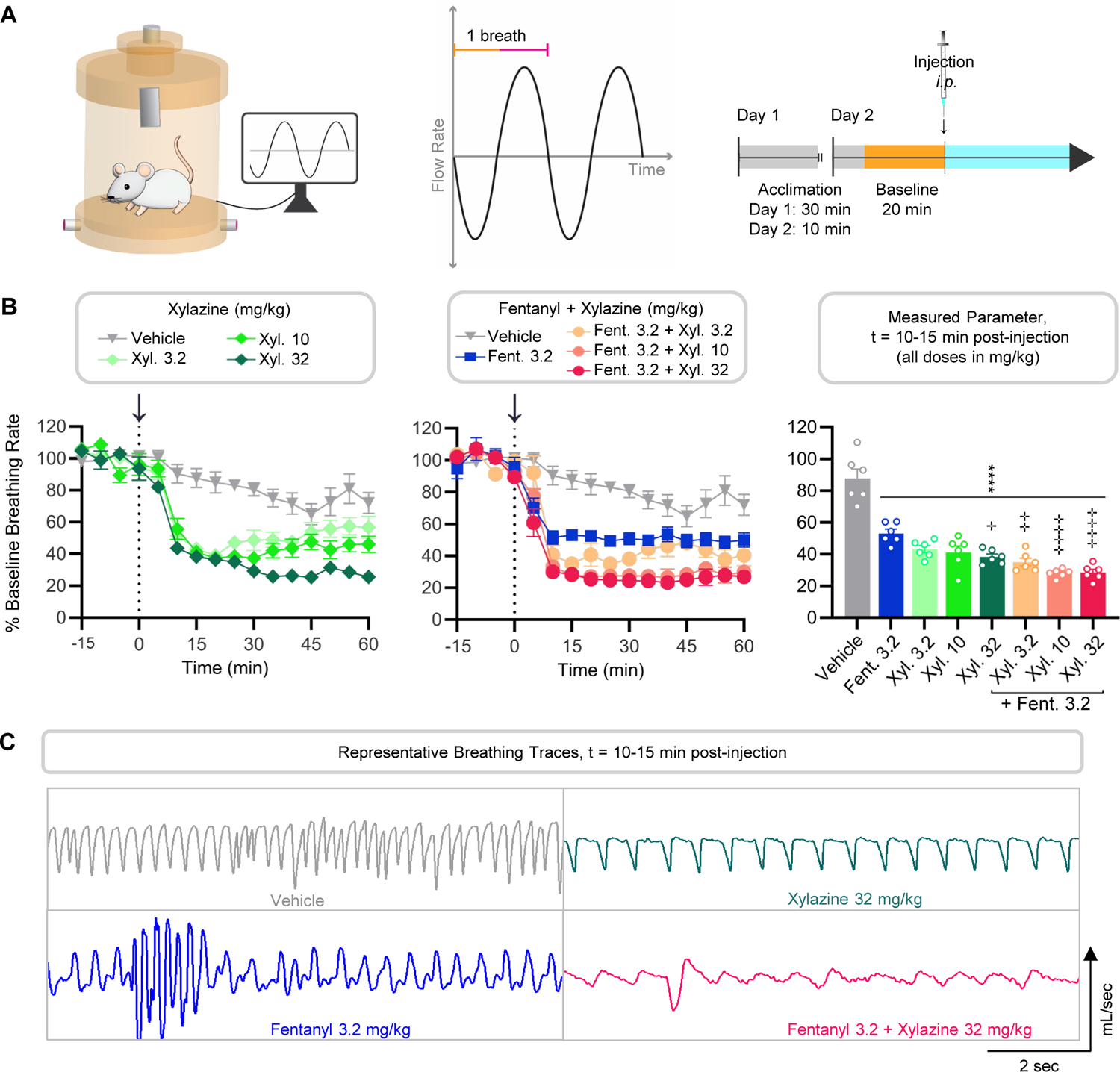
Xylazine reduces breathing rate to a greater degree than fentanyl and drives the effect when the two drugs are combined. **A)** Graphical representation of study design. Breathing rate of male and female CD-1 mice (n=6 per group) was recorded using vivoFlow^®^ whole-body plethysmography (WBP) chambers supplied with air according to the timeline. **B)** Arrows and dotted lines indicate *i.p.* injection (t=0). Breathing rate was compared among all 9 groups via two-way ANOVA (t=0-60 min; main effect of treatment p<0.0001). Average values by treatment group from t=10-15 min post-injection were compared via one-way ANOVA (p<0.0001) with Tukey’s post-hoc test (p-values displayed). **C)** Representative breathing traces from selected treatment groups from t=10-15 min post-injection. ****p<0.0001 vs. vehicle; **^⊹^**p<0.05, **^⊹⊹^**p<0.01, **^⊹⊹⊹⊹^**p<0.0001 vs. fentanyl alone

### Xylazine exacerbates fentanyl-induced changes in inspiration

As opioids primarily inhibit brainstem regions that control the process of inspiration [24–27], we next evaluated how xylazine alters fentanyl-induced changes in inspiration time (Ti) and peak inspiratory flow (PIF), as depicted in **Fig. 2A**. Relative to vehicle, xylazine did not affect inspiration time at any dose tested (**Fig. 2B**). As expected, fentanyl alone and in combination with xylazine (3.2, 10, or 32 mg/kg) increased inspiration time compared to vehicle (p<0.0001, t=10-15 min post-injection; **Fig 2B** bar graph). Only fentanyl combined with 10 mg/kg xylazine produced a significant increase in inspiration time relative to fentanyl alone (p<0.05). Similarly, xylazine did not affect peak inspiratory flow (**Fig. 2C**), while fentanyl alone and in combination with xylazine decreased peak inspiratory flow relative to vehicle (p<0.05 and p<0.0001, t=10-15 min post-injection; **Fig 2C** bar graph). The combination of fentanyl and xylazine (3.2, 10, or 32 mg/kg) reduced peak inspiratory flow below that of fentanyl at all doses (p<0.01, p<0.01, and p<0.05, respectively), indicating a supra-additive effect on peak inspiratory flow (**Fig. 2C**). Taken together, these results show that xylazine lacks an independent effect on inspiration but enhances fentanyl-induced changes in inspiration.

**Figure 2.**
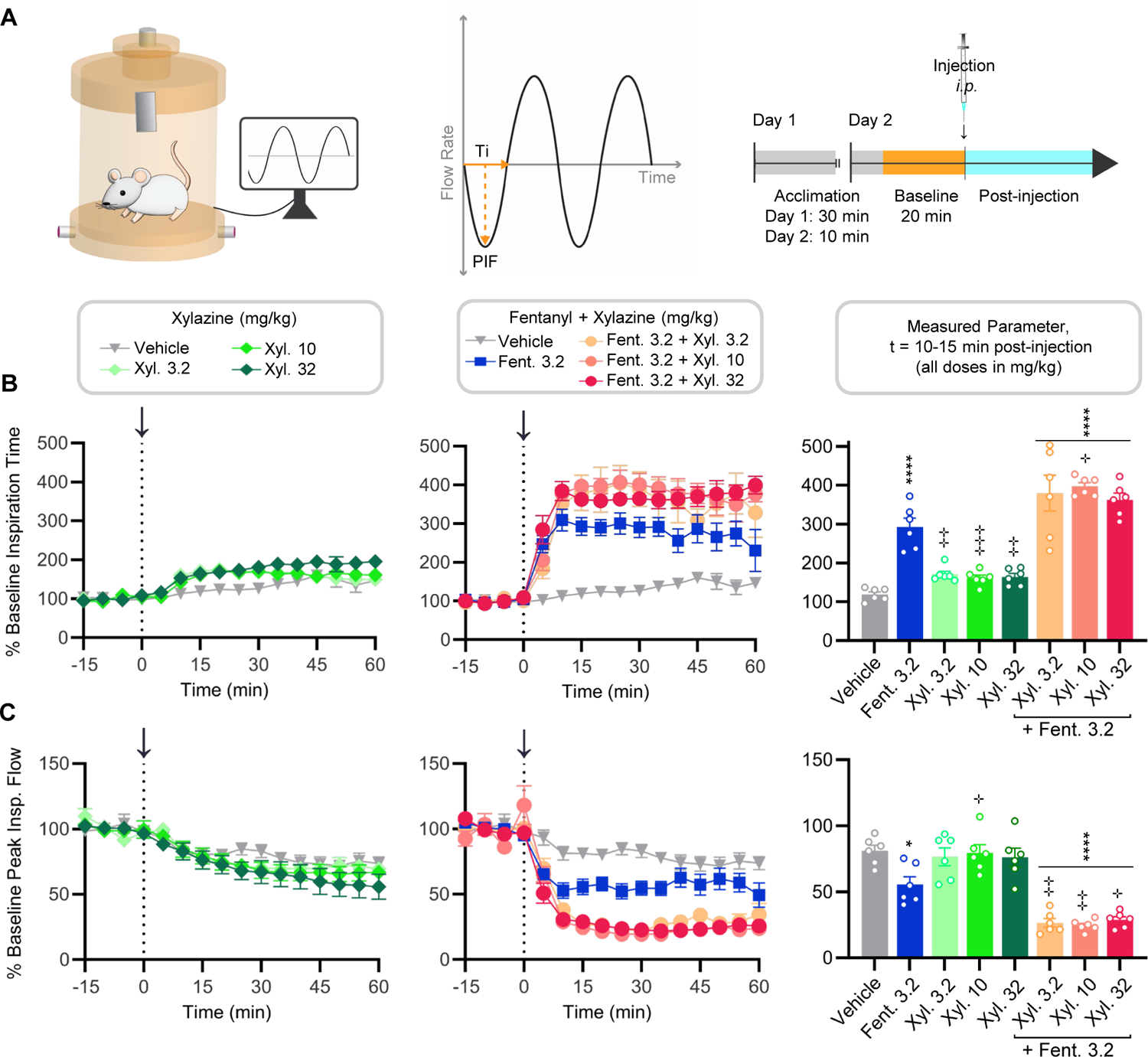
Xylazine exacerbates fentanyl-induced changes in inspiration. **A)** Graphical representation of study design. Inspiratory parameters of male and female CD-1 mice (n=6 per group) were recorded using vivoFlow^®^ WBP chambers supplied with air according to the timeline. **B-C)** Arrows and dotted lines indicate *i.p.* injection point. Inspiratory parameters were compared independently among all 9 groups via two-way ANOVA (t=0-60 min; main effect of treatment p<0.0001 for both parameters). Average values by treatment group from t=10-15 min post-injection were compared via one-way ANOVA (p<0.0001 for both parameters) with Tukey’s post-hoc test (p-values displayed). **B)** Time course of inspiration time (Ti) from 0-60 min by treatment group with a bar graph displaying the group values by from t=10-15 min (average ± S.E.M.). **C)** Time course of peak inspiratory flow (PIF) from 0-60 min by treatment group with a bar graph displaying the group values from t=10-15 min (average ± S.E.M.). *p<0.05, ****p<0.0001 vs. vehicle; **^⊹^**p<0.05, **^⊹⊹^**p<0.01, **^⊹⊹⊹^**p≤0.001 vs. fentanyl alone

### Xylazine produces robust effects on expiration that differ from those induced by fentanyl

The effects of fentanyl and xylazine on expiration time (Te) and peak expiratory flow (PEF), illustrated in **Fig. 3A**, were also considered. We found that xylazine drastically increased expiration time while fentanyl had no effect (**Fig. 3B**). From 10-15 min post-injection, all doses of xylazine produced significant increases in expiration time relative to vehicle (3.2 mg/kg: p<0.01; 10 mg/kg: p<0.001; 32 mg/kg: p<0.01), as did fentanyl in combination with 10 or 32 mg/kg xylazine (p<0.01 and p<0.0001, respectively). Fentanyl did not increase the effect of any dose of xylazine on expiration time. The large effect of xylazine on expiration time accounts for the xylazine-driven reduction in breathing rate observed in **Fig. 1B**. Xylazine decreased peak expiratory flow relative to control (**Fig. 3C**) at all doses tested (p<0.01). Interestingly, fentanyl significantly increased peak expiratory flow relative to control (p<0.0001). When fentanyl was combined with xylazine, peak expiratory flow was significantly decreased relative to fentanyl alone (p<0.0001, t=10-15 min post-injection) to levels comparable to xylazine alone. This suggests that xylazine reverses the fentanyl-specific increase in peak expiratory flow. We observed similar findings for expiratory flow 50 (EF_50_), the flow rate when half of the volume of air in the lungs has been expired (**Fig. 3D**). Fentanyl increased EF_50_ (p<0.0001) while xylazine at 3.2 (p<0.05 for t=0-15 min; p<0.001 for t=0-60 min) and 32 mg/kg (p<0.0001 for t=0-15 min and t=0-60 min) decreased this parameter relative to control. As with peak expiratory flow, the combination of fentanyl and either 3.2 or 32 mg/kg xylazine yielded a significant reduction in EF_50_ relative to fentanyl alone (p<0.0001 for all at t=0-15 min and t=0-60 min) to levels comparable to control from 0-15 min post-injection. Only the 32 mg/kg dose of xylazine produced a significantly lower EF_50_ alone versus in combination with fentanyl from t=0-15 min (p<0.01) and t=0-60 min (p<0.05). In summary, xylazine greatly affects the process of expiration and reverses fentanyl-induced increases in expiratory flow.

**Figure 3.**
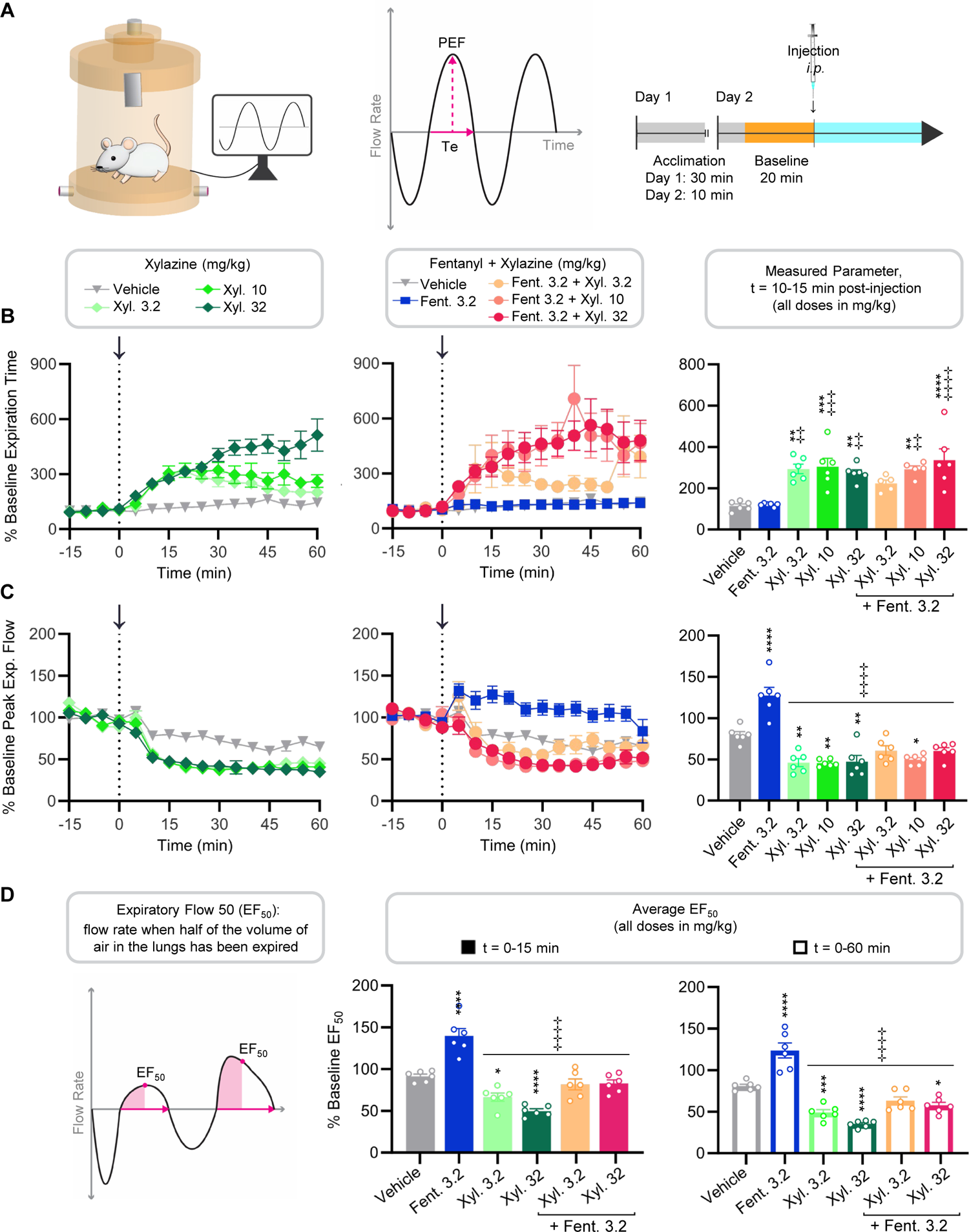
Xylazine produces robust effects on expiration that differ from those induced by fentanyl. **A)** Graphical representation of study design. Expiratory parameters of male and female CD-1 mice (n=6 per group) were recorded using vivoFlow^®^ WBP chambers supplied with air according to the timeline. **B-C)** Arrows and dotted lines indicate *i.p.* injection point. Expiratory parameters were compared independently among all 9 groups via two-way ANOVA (t=0-60 min; main effect of treatment p<0.0001 for both parameters). Average values by treatment group from t=10-15 min post-injection were compared via one-way ANOVA (p<0.0001 for both parameters) with Tukey’s post-hoc test (p-values displayed). **B)** Time course of expiration time (Te) from 0-60 min by treatment group with a bar graph displaying the group values from t=10-15 min (average ± S.E.M.). **C)** Time course of peak expiratory flow (PEF) from 0-60 min by treatment group with a bar graph displaying the group values from t=10-15 min (average ± S.E.M.). **D)** Expiratory flow 50 (EF_50_; average ± S.E.M.) values by treatment group from t=0-15 min (filled bar graph) or t=0-60 min post-injection (open bar graph) were compared via one-way ANOVA (p<0.0001 for both time ranges) with Tukey’s post-hoc test (p-values displayed). *p<0.05, **p<0.01, ***p<0.001, ****p<0.0001 vs. vehicle; **^⊹⊹^**p<0.01, **^⊹⊹⊹^**p<0.001, **^⊹⊹⊹⊹^**p<0.0001 vs. fentanyl alone

### Fentanyl but not xylazine decreases blood oxygen saturation and both drugs lower heart rate

We also examined how fentanyl and xylazine affect blood oxygen levels using pulse oximetry (as depicted in **Fig. 4A**), since depleted blood oxygen saturation is what ultimately leads to fatality in an overdose. Percent blood oxygen saturation is shown for xylazine, fentanyl, and fentanyl combined with xylazine (**Fig. 4B**). Xylazine had no impact on blood oxygen saturation alone, while fentanyl or fentanyl plus xylazine reduced blood oxygen levels compared to vehicle (p<0.0001; **Fig. 4B**). There were no differences in the maximal reduction in blood oxygen saturation (t=10-15 min) between fentanyl and either combination of fentanyl plus xylazine. We also compared the area under the curve from t=0-60 min (**Fig. 4B** bar graph). Similarly, only treatments containing fentanyl, alone or in combination with xylazine, decreased blood oxygen saturation compared to vehicle (p<0.0001). These results show that doses of 3.2-32 mg/kg xylazine do not contribute to blood oxygen saturation loss as measured by pulse oximetry.

**Figure 4.**
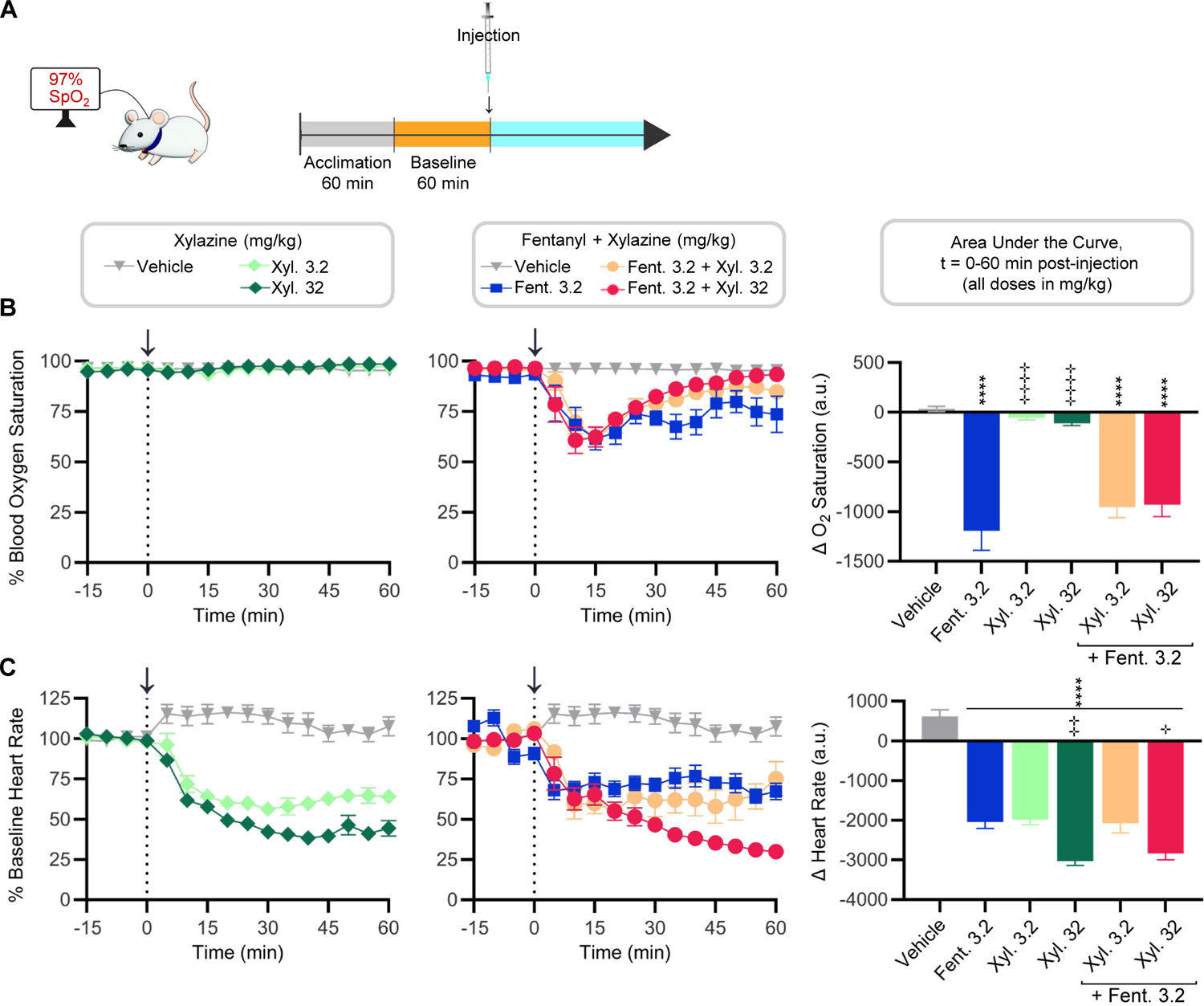
Fentanyl but not xylazine decreases blood oxygen saturation and both drugs lower heart rate. **A)** Graphical representation of study design. Blood oxygen saturation and heart rate of male and female CD-1 mice (n=6 per group) were measured using the MouseOx^®^ pulse oximetry system according to the timeline shown. **B-C)** Arrows and dotted lines indicate *i.p.* injection point. Area under the curve was compared independently among all 6 groups via two-way ANOVA (t=0-60 min; main effect of treatment p<0.0001 for both parameters). Area under the curve values by treatment group from t=0-60 min post-injection were compared via one-way ANOVA (p<0.0001 for both parameters) with Tukey’s post-hoc test (p-values displayed in bar graph). **B)** Time course of blood oxygen saturation (%) by treatment group with a bar graph displaying area under the curve values from t=0-60 min post-injection (average ± S.E.M.). **C)** Time course of heart rate by treatment group with a bar graph displaying area under the curve values from t=0-60 min post-injection (average ± S.E.M.). ****p<0.0001 vs. vehicle; **^⊹^**p<0.05, **^⊹⊹^**p<0.01, **^⊹⊹⊹⊹^**p<0.0001 vs. fentanyl alone

As xylazine is known to induce severe bradycardia in humans, we also evaluated heart rate. Xylazine, fentanyl, and both combinations of fentanyl plus xylazine (3.2 or 32 mg/kg) significantly decreased heart rate relative to vehicle at t=10-15 min (p<0.0001). We also calculated the area under the curve for each treatment (**Fig. 4C** bar graph). Again, each treatment significantly reduced heart rate relative to vehicle (p<0.0001). There was no difference in heart rate reduction between xylazine alone or in combination with fentanyl at either dose, indicating a sub-additive effect. The dose of 32 mg/kg xylazine significantly decreased heart rate, alone or in combination with fentanyl, relative to fentanyl alone (p<0.01 and p<0.05). These results indicate that high doses of xylazine – alone or in combination with fentanyl – decrease heart rate more than fentanyl over time.

### Fentanyl induces severe apneas that correlate with loss in oxygen saturation

Given that fentanyl in combination with xylazine reduced breathing rate (**Fig. 1B**), and peak inspiratory (**Fig. 2C**) and expiratory (**Fig. 3C**) flow compared to fentanyl alone, we were surprised that xylazine did not worsen fentanyl-induced changes in oxygen saturation (**Fig. 4B**). It is known that apneas are associated with reduced blood oxygen saturation [28,29], thus we next evaluated whether apneas could explain this discrepancy (**Fig 5**). As depicted in **Fig. 5A**, an apnea was defined as a breath with expiration time (Te) longer than twice the average post-injection expiration time calculated for each individual mouse: *Apnea == Te > 2[Avg. Te post-injection]*. A severe apnea was classified as a breath in which the expiration time exceeded twice the average breath length (expiration and inspiration time) post-injection for each individual mouse: *Severe Apnea == Te > 2[Avg. Ti + Te post-injection]*. The total number of apneas (**Fig. 5B**) and severe apneas (**Fig. 5C**) are shown for the first t=0-15 min post-injection, during which oxygen saturation is maximally decreased. As shown in **Fig. 5B**, fentanyl alone or in combination with 3.2 mg/kg xylazine induced a significant number of total apneas compared to control (p<0.0001 and p<0.01, respectively). A significant correlation between the number of total apneas and blood oxygen saturation in the first 15 min post-injection was found (p<0.05, R^2^=0.7088; **Fig. 5D**). By restricting analysis to severe apneas only, we found that fentanyl alone (p<0.01) or in combination with xylazine (p<0.05) induced significantly more apneas than vehicle (**Fig. 5C**). There was a strong correlation between the number of severe apneas and oxygen saturation loss (p<0.0001, R^2^=0.9996; **Fig. 5E**). Taken together, the fentanyl-specific decrease in blood oxygen saturation may be explained by the greater frequency of severe apneas induced by fentanyl during the first 15 min post-injection, during which time oxygen saturation is maximally reduced.

**Figure 5.**
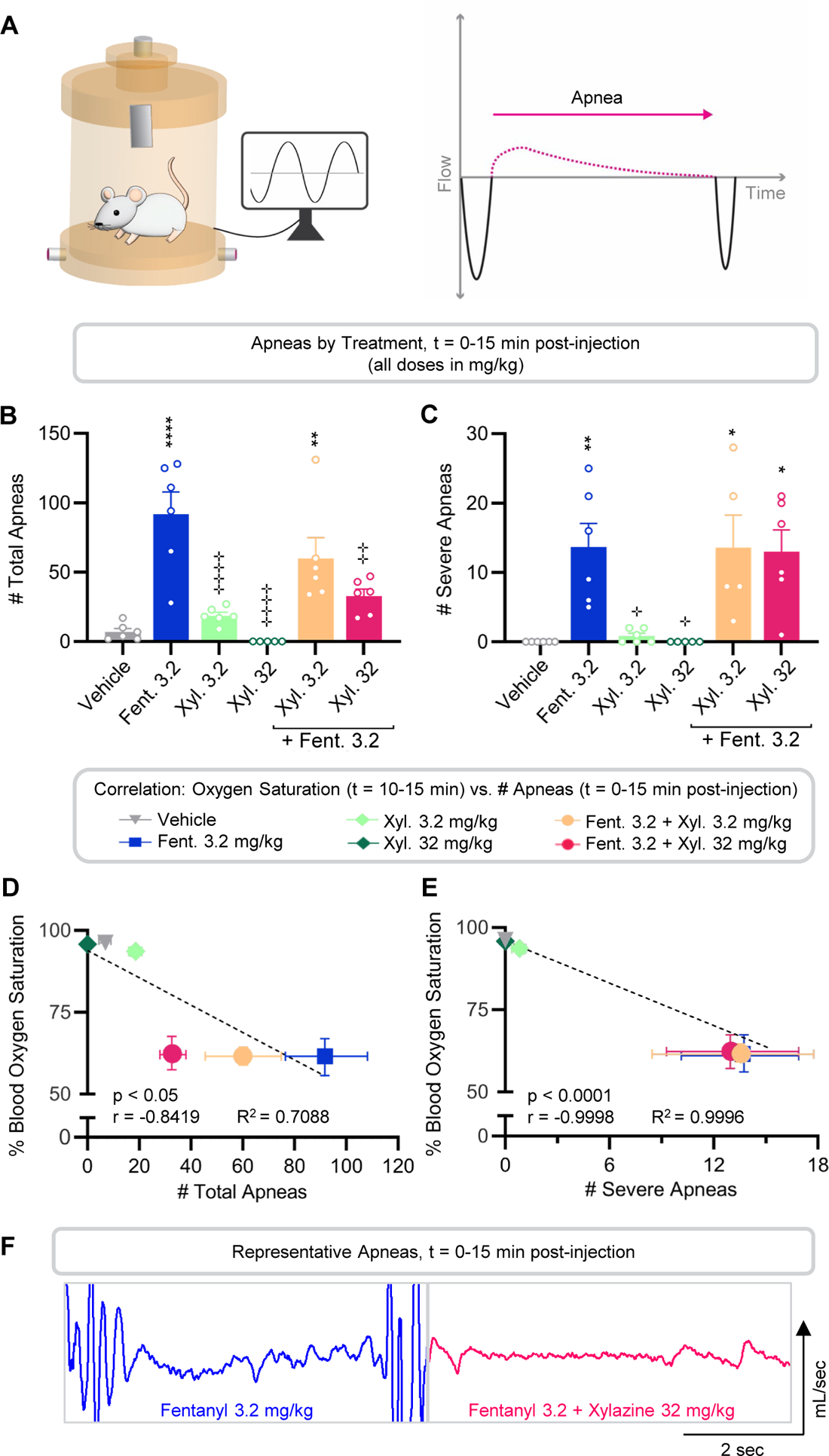
Fentanyl induces severe apneas that correlate with loss in oxygen saturation. **A)** Graphical representation of study design. Frequency of apneas in male and female CD-1 mice (n=6 per group) was determined using vivoFlow^®^ WBP chambers supplied with air. **B)** Number of total apneas from t=0-15 min post-injection by treatment group (average ± S.E.M.) were compared via one-way ANOVA (p<0.0001) with Tukey’s post-hoc test (p-values displayed). *Apnea = Te > 2[Avg.Te post-injection]*. **C)** Number of severe apneas from t=0-15 min post-injection by treatment group (average ± S.E.M.) were compared via one-way ANOVA (p<0.001) with Tukey’s post-hoc test (p-values displayed). *Severe Apnea* > 2[Avg.Te post-injection]. **D)** Pearson correlation (p<0.05) between % oxygen saturation (t=10-15 min post-injection) and # total apneas (t=0-15 min post-injection). Each data point is the average ± S.E.M. per group. Linear equation: *y = −0.44x + 93.97*. **E)** Pearson correlation (p<0.0001) between % oxygen saturation (t=10-15 min post-injection) and # severe apneas (t=0-15 min post-injection). Linear equation: *y* = −2.54*x* + 95.94. **F)** Representative apnea from t=10-15 min post-injection. *p<0.05, **p<0.01, ****p<0.0001 vs. vehicle; **^⊹^**p<0.05, **^⊹⊹^**p<0.01, **^⊹⊹⊹⊹^**p<0.0001 vs. fentanyl alone 2 sec

## DISCUSSION

In mammals, breathing rate is governed by active inspiration and passive expiration. Herein, we show that xylazine produces a greater reduction in breathing rate than fentanyl. In contrast to fentanyl, which increases inspiration time, xylazine reduces breathing rate by extending expiration time. Although xylazine did not produce any changes in inspiration alone, it did exacerbate the effects of fentanyl. Alone, fentanyl but not xylazine reduced blood oxygen saturation at the doses tested, and xylazine did not further reduce blood oxygen saturation when combined with fentanyl. The reduction in blood oxygen caused by fentanyl strongly correlated with the frequency of severe apneas; xylazine did not cause apneas or exacerbate the effects of fentanyl. Finally, both fentanyl and xylazine induced bradycardia, with higher doses of xylazine driving the effect when the two drugs were combined. Altogether, these results show that xylazine exacerbates fentanyl-induced depression of respiration and heart rate.

Our results showing that fentanyl primarily affected the process of inspiration are consistent with a body of prior literature and reflect opioid inhibition of neurons in the pre-Bötzinger complex and Kölliker-Fuse nucleus of the brainstem [2,4,25,26,30–32]. Fentanyl increases inspiration time, thereby lowering breathing rate, and also reduces peak inspiratory flow. When combined with fentanyl, all doses of xylazine administered further reduced peak inspiratory flow in what appears to be a supra-additive manner. Considering xylazine did not change inspiratory flow when given alone, the supra-additive effect when combined with fentanyl may reflect the greater reliance on brainstem respiratory rhythm generation – which is inhibited by opioids – when sleeping [33] or sedated by drugs like xylazine.

During passive expiration, neurons in the ventral rostral group of the brainstem fire intrinsically as part of the central pattern generator during passive expiration [34,35]. Xylazine could inhibit expiration through α2 adrenergic receptor (α2AR) autoreceptor activation in these neurons [36], or in chemosensory neurons within the locus coeruleus [37] or nucleus tractus solitarius [38]. Xylazine also reduced peak expiratory flow and EF_50_, while fentanyl increased these parameters at a dose that did not increase locomotion. In rodents, changes in EF_50_ have been observed in response to respiratory challenges such as SARS-CoV-2 infection [39] and allergic asthma [40]. In line with our results, fentanyl was shown to induce a prolonged increase in EF_50_ following fentanyl administration in rats [41], yet this finding has not been consistent across all studies [42]. It’s possible that the opposing effects of fentanyl and xylazine on EF_50_ reflect the respiratory muscle rigidity caused by fentanyl [3] versus the muscle relaxant effects of xylazine [43], but more studies are needed to better understand this metric in the context of opioid use.

The correlation between the fentanyl-induced reduction in blood oxygen saturation and number of apneas is consistent with human data that show a greater decrease in blood oxygen levels associated with severity of obstructive sleep apnea [28,29]. Consistent with our findings using pulse oximetry, anesthetic doses of xylazine were observed to decrease breathing rate with little effect on peripheral blood oxygen saturation [44]. Moreover, it was shown via electrochemistry that xylazine did not worsen the maximum level of hypoxia of fentanyl in the nucleus accumbens (NAc) of rats [18,45]. However, these studies also found that xylazine on its own decreased oxygenation of the NAc [18,45] whereas the current experiments show no effect on blood oxygen saturation, potentially reflecting different sensitivities of the techniques used. It is important to mention that mice have a much higher specific metabolic rate than humans [46,47], which allows them to better adapt to changes in blood oxygen saturation.

As a sedative, xylazine is known to decrease metabolic rate [48,49], which decreases breathing rate. It’s therefore possible that xylazine does not directly inhibit breathing circuits but instead reduces breathing rate by decreasing metabolic rate. This may account for the lack of effect on blood oxygen saturation: the lower breathing rate induced by xylazine may be sufficient under conditions of decreased metabolic demand. Separately, high CO_2_ conditions (e.g., 5%) mimic increased metabolic demand [50,51] and are utilized in many opioid-induced respiratory depression studies to increase ventilatory drive [2,26]. This is a confounding variable because hypercapnia-driven breathing is mechanistically distinct and opioids inhibit the ventilatory response to hypercapnia, which can magnify a drug’s perceived effect. It is worth noting that we used air in these studies to better reflect the context of human overdose and avoid these confounds.

Finally, our data showing that xylazine alone or in combination with fentanyl induced bradycardia to a greater degree than fentanyl alone are consistent with previous literature reporting decreased heart rate and hypotension as clinical symptoms of fentanyl-xylazine overdose [52]. We have established that fentanyl lowers blood oxygen levels and xylazine decreases heart rate, reducing oxygen supply to critical tissues. In a fatal overdose, inadequate oxygenation of the brain and heart cause brain damage and eventual cardiac arrest [24,53]. Thus, the xylazine-driven changes in the cardiovascular system coupled with the low blood oxygen levels induced by fentanyl could greatly increase the risk of overdose in humans.

Altogether, we show that xylazine worsens fentanyl-induced respiratory depression via direct effects on expiration, as well as exacerbating fentanyl’s effects on inspiration. Despite the combination of fentanyl and xylazine producing a severely lowered breathing rate than fentanyl alone, this did not translate to a further reduction in blood oxygen saturation. Instead, blood oxygen depletion correlated with the number of apneas. Additionally, xylazine reduced heart rate more than fentanyl and was primarily responsible for this effect when combined. Thus, xylazine produces independent effects on respiration and heart rate while also exacerbating those produced by fentanyl. These results provide preclinical insight into the mechanisms underlying xylazine’s contribution to overdoses involving opioids in humans. Future work should delineate how these effects manifest in the overall human population with regard to risk of fatal overdose.

## AUTHOR CONTRIBUTIONS

Catherine Demery-Poulos: Conceptualization, Experimental Design, Data Collection and Analysis, Writing Sierra C. Moore: Data Collection and Analysis Erica S. Levitt: Experimental Design, Editing Jessica P. Anand: Conceptualization, Experimental Design, Funding, Project Administration, Editing John R. Traynor: Conceptualization, Experimental Design, Funding, Project Administration, Editing

